# Loss of H3K9 trimethylation leads to premature aging

**DOI:** 10.1101/2024.07.24.604929

**Authors:** Calida Mrabti, Na Yang, Gabriela Desdín-Micó, Alejandro Alonso-Calleja, Alba Vílchez-Acosta, Sara Pico, Alberto Parras, Yulan Piao, Lucas Schoenfeldt, Siyuan Luo, Amin Haghani, Robert Brooke, María del Carmen Maza, Clémence Branchina, Céline Yacoub Maroun, Ferdinand von Meyenn, Olaia Naveiras, Steve Horvath, Payel Sen, Alejandro Ocampo

**Affiliations:** Department of Biomedical Sciences, Faculty of Biology and Medicine, University of Lausanne, Lausanne, Vaud, Switzerland; Laboratory of Genetics and Genomics, National Institute on Aging, NIH, Baltimore, MD 21224, USA; Laboratory of Regenerative Hematopoiesis, Department of Biomedical Sciences, University of Lausanne, Switzerland; Laboratory of Metabolic Signaling, Institute of Bioengineering, Ecole Polytechnique Fédérale de Lausanne, Lausanne, Switzerland; EPITERNA SA, Epalinges, Switzerland; Departement of Health Sciences and Technology, ETH Zurich, Zurich; Altos Labs, San Diego, CA, USA; Epigenetic Clock Development, Foundation, Torrance, California, USA; Human Genetics, David Geffen School of Medicine, University of California, Los Angeles, CA, USA

**Keywords:** H3K9me3, aging, epigenetics, chromatin

## Abstract

Aging is the major risk factor for most human diseases and represents a major socio-economical challenge for modern societies. Despite its importance, the process of aging remains poorly understood. Epigenetic dysregulation has been proposed as a key driver of the aging process. Modifications in transcriptional networks and chromatin structure might be central to age-related functional decline. A prevalent feature described during aging is the overall reduction in heterochromatin, specifically marked by the loss of repressive histone modification, Histone 3 lysine 9 trimethylation (H3K9me3). However, the role of H3K9me3 in aging, especially in mammals, remains unclear. Here we show using a novel mouse strain, (TKOc), carrying a triple knockout of three methyltransferases responsible for H3K9me3 deposition, that the inducible loss of H3K9me3 in adulthood results in premature aging. TKOc mice exhibit reduced lifespan, lower body weight, increased frailty index, multi-organ degeneration, transcriptional changes with significant upregulation of transposable elements, and accelerated epigenetic age. Our data strongly supports the concept that the loss of epigenetic information directly drives the aging process. These findings reveal the importance of epigenetic regulation in aging and suggest that interventions targeting epigenetic modifications could potentially slow down or reverse age-related decline. Understanding the molecular mechanisms underlying the process of aging will be crucial for developing novel therapeutic strategies that can delay the onset of age-associated diseases and preserve human health at old age specially in rapidly aging societies.

## INTRODUCTION

In the last few years, the field of epigenetics has gained prominence, holding significant implications for various aspects of human health. During aging, epigenetic dysregulation is observed leading to changes in gene expression (Sen *et al*., 2016). In this context, aging is associated with a global loss and a local increase in DNA methylation. In this line, it is relevant to highlight that novel epigenetic clocks based on these age-associated changes in DNA methylation have been developed by multiple groups (Horvath, 2013; Field *et al*., 2018). These clocks provide a valuable tool for understanding the complex interplay between epigenetic modifications and the aging process. In addition to changes in DNA methylation, alterations in the tri-methylation of histone 3 lysine 9 (H3K9me3) associated with repressive heterochromatin (Peters *et al*., 2003; Rea *et al*., 2000; Montavon *et al*., 2021) occur during aging in model organisms (Booth and Brunet, 2016), in human samples of individuals at an advanced age, and in patients suffering from premature aging syndromes including Hutchinson–Gilford progeria syndrome and Werner syndrome (Scaffidi and Misteli, 2006; Shumaker *et al*., 2006; Zhang *et al*., 2015b). In addition, the levels of the H3K9me3 methyltransferase Suv39h1 and the heterochromatin protein 1 (HP1) decrease during normal aging due to alterations in other hallmarks of aging such as DNA damage, telomere shortening, and mitochondrial dysfunction (Zhang *et al*., 2015a).

Importantly, the decrease in heterochromatin results in increased transcriptional activity in non-coding regions of the genome, including repetitive regions containing transposable elements (TEs), which are generally repressed by H3K9me3 (De Cecco, Criscione, Peckham, *et al*., 2013; De Cecco, Criscione, Peterson, *et al*., 2013; Wood and Helfand, 2013; He *et al*., 2019; Gorbunova *et al*., 2021). TEs can be broadly divided into DNA transposons or retrotransposons, which make cDNA copies through reverse transcription. Retrotransposons are further classified as long terminal repeats (LTR)-containing endogenous retroviruses (ERVs) or non-LTR retrotransposons such as long interspersed nuclear elements (LINEs) or short interspersed nuclear elements (SINEs). With age, transcriptional activation of retrotransposons can have detrimental consequences such as activation of innate immunity, DNA damage, and various diseases such as cancer and autoimmune diseases (Gorbunova *et al*., 2021; De Cecco *et al*., 2019). Based on these evidences, the global loss of H3K9me3 observed in multiple species during aging has led to the "heterochromatin loss theory of aging" (Villeponteau, 1997; Tsurumi and Li, 2012). Although these observations suggest a central role of the age-associated loss of heterochromatin and H3K9me3 as drivers of the aging process, this role has not been demonstrated yet in mammals, where current data only shows a correlation between loss of H3K9me3 and aging.

To investigate the potential role of epigenetic dysregulation in the aging process, we aimed to create an experimental mouse model featuring the inducible loss of H3K9me3 during adulthood. To achieve this goal, we selected a genetic approach involving the inducible knockout of the three methyltransferases, Suv39h1, Suv39h2, and Setdb1, known for establishing the H3K9me3 mark. Due to the previously demonstrated essential role of H3K9me3 during development, we induced H3K9me3 loss in adult mice (Peters *et al*., 2001a; Dodge *et al*., 2004; Tachibana *et al*., 2007; Nicetto *et al*., 2019a). Importantly, loss of H3K9me3 resulted in reduced lifespan and was associated with multiple age-associated phenotypic alterations, indicating that the loss of the epigenetic mark H3K9me3 can drive the aging process.

## RESULTS

### Novel mouse model for the inducible loss of H3K9me3 at adult stage

The generation of a triple knock out strain (TKO) for the three H3K9me3 methyltransferases Suv39h1, Suv39h2 and, Setdb1 has been previously reported for investigating the role of H3K9me3 during development (Nicetto *et al*., 2019a). In this study, the disruption of these three methyltransferases during development demonstrated the crucial role of H3K9me3 during the initiation of organogenesis, as well as in the preservation of lineage fidelity (Nicetto *et al*., 2019a).

In order to investigate the role of H3K9me3 during aging and bypass the problems associated with the loss of H3K9me3 during development (Peters *et al*., 2001b; Dodge *et al*., 2004; Tachibana *et al*., 2007; Nicetto *et al*., 2019a), we designed a transgenic strategy to conditionally induce H3K9me3 depletion at adult stage by using a tamoxifen-inducible Cre-mediated recombination system. To do so, TKO mice (Nicetto *et al*., 2019a) were crossed with the constitutive CAG-CreER mouse (Hayashi and McMahon, 2002). This new strain, TKOCAGCre, allows the activation of whole-body Cre recombinase upon tamoxifen administration leading to the deletion of Setdb1 and Suv39h1 while Suv39h2 is constitutively knocked out (Figure 1A; Figure S1A).

**Figure 1.**
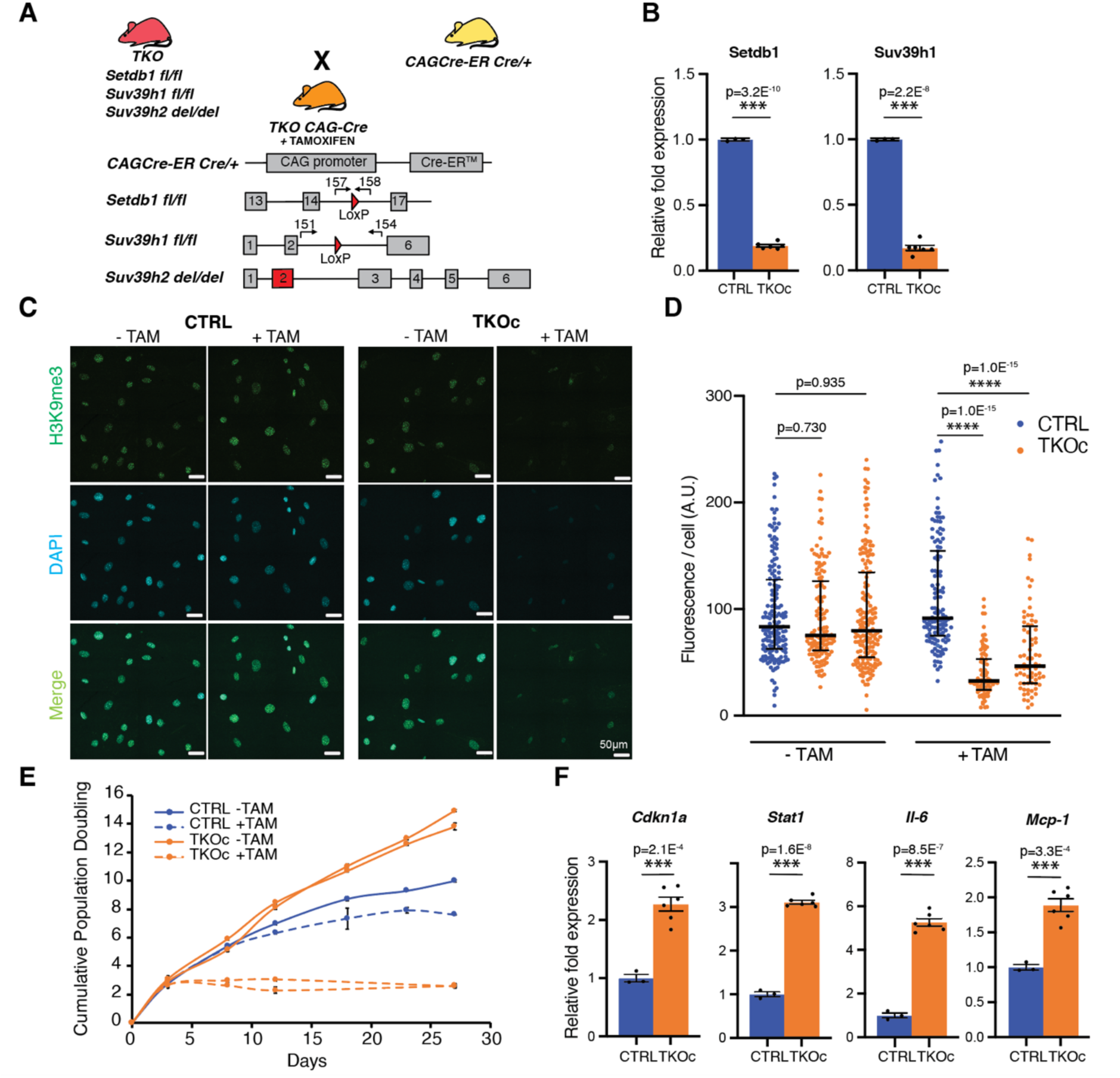
Loss of H3K9me3 *in-vitro* induces cell cycle arrest and cellular senescence. **(A)** Schematic representation of the genetic strategy to generate the quadruple transgenic TKOCAGCre mouse strain carrying the insertion of CAG-CreER (Chr.3) for the tamoxifen-inducible CRE-mediated recombination system (Loxp sites), Setdb1 gene insertion of LoxP sites in the intron 14 and 16 (Chr.3), Suv39h1 gene insertion of LoxP sites in the intron 2 and 5 (Chr.X), Suv39h2 gene deletion at Exon2 (Chr. 10). **(B)** Setdb1 and Suv39h1 mRNA levels in CTRL and TKOc tail-tip fibroblasts after 6 days of 4-OH Tamoxifen treatment. (n = 1 to 2, with 3 technical replicate). **(C)** Immunofluorescence of H3K9me3 in CTRL and TKOc tail-tip fibroblasts upon tamoxifen treatment for 6 days. Scale bar, 50 μm. **(D)** Quantification of H3K9me3 positive cells in CTRL and TKOc tail-tip fibroblasts upon tamoxifen treatment. *p < 0.05; **p < 0.01; ***p < 0.001; ****p < 0.0001 according to one-way ANOVA data are median with interquartile range. **(E)** Western blot of H3K9me3 in CTRL and TKOc tail-tip fibroblasts with or without TAM treatment. H3K9me3 quantification in CTRL and TKOc after treatment. **(F)** Cumulative population doubling curves of CTRL and TKOc tail-tip fibroblasts with or without TAM treatment. Population doublings were calculated by the formula log [(number of cells harvested)/(number of cells seeded)]/ log2. **(G)** Relative mRNA levels of genes related to senescence markers and senescence-associated secretory phenotype in CTRL and TKOc tail-tip fibroblasts after TAM treatment. (n = 1 to 2, with 3 technical replicate). Data are mean ± SEM. Statistical significance was assessed by Two-tailed Student’s t test. *p < 0.05; **p < 0.01; ***p < 0.001.

First, to validate the genetic strategy, tail tip fibroblast (TTFs) from TKO CAG-Cre mice both positive (TKOc) and negative (CTRL) for CAG-Cre were isolated and cultured. After 2 days in culture, TTFs were treated with 4-hydroxy tamoxifen (4-OHT) for 6 days and mRNA levels of methyltransferases were analyzed. As expected, we observed a downregulation of Setdb1 and Suv39h1 (Figure 1B). In addition, a decrease in the protein levels of H3K9me3 was detected by immunofluorescence and Western blot upon 4-OHT treatment of TKOc TTFs compared to CTRLs (Figure 1C-E). Interestingly, loss of H3K9me3 caused a decrease in cell proliferation in TKOc TTFs upon 4-OHT treatment (Figure 1F).

Next, to determine whether cell cycle arrest was due to cellular senescence, we measured the expression of markers of cellular senescence and senescence-associated secretory phenotype (SASP) in TKOc cells. Interestingly, we observed an increase in cellular senescence in TKOc treated cells with upregulation of *Cdkn1a* (cyclin dependent kinase inhibitor 1A) (Campisi and d’Adda di Fagagna, 2007) and *Stat1* (signal transducer and activator of transcription) mRNA (Novakova *et al*., 2010). In addition, the expression of genes associated with SASP, including *Il-6* (interleukin-6) and *Mcp-1* (monocyte chemoattractant protein-1), were upregulated in TKOc cells due to the loss of H3K9me3 (Figure 1G). In addition, a significant increase in SA-beta-galactosidase activity was detected upon 4-OHT treatment of TKOc TTFs compared to CTRLs (Figure S1C). Together, these data indicate that our genetic strategy allows the inducible depletion of H3K9me3 leading to cell cycle arrest and cellular senescence.

### Loss of H3K9me3 leads to premature aging

To investigate whether H3K9me3 reduction in adult mice could drive premature aging, we designed an induction protocol were 3-month-old mice were treated with 5 consecutive daily intraperitoneal injections of tamoxifen (TAM) (Figure S2A). First, we validated the system in vivo by confirming the recombination of Setdb1 and Suv39h1 in DNA isolated from peripheral blood of TKOc and control three months after treatment with TAM for 5 days (Figure S2B). TKOc mice started to exhibit a moderate change in their physical appearance at 6 months of age, prompting us to perform health status and behavioral assessments. Towards this goal, we used a composite frailty index (FI) score to assess health measures including body and coat condition, kyphosis, cataract, and tail stiffening (Whitehead *et al*., 2014). TKOc mice presented higher FI scores than CTRL mice (Figure S2C). In addition, we monitored changes in body weight and did not detect remarkable changes between treated TKOc and their CTRL groups over a period of 42 weeks (Figure S2D). On the other hand, analysis of hematological parameters by complete blood count (CBC) showed a significant decrease in red blood cell (RBC) and hemoglobin (HGB) levels, indicating signs of anemia in TKOc treated mice (Figure S2E). Moreover, analysis of activity by open field test indicated that TKOc mice exhibited hypoactivity and slower movements together with an increase in peripheral distance travelled compared to the CTRL group (Figure S2F). Contrary, grip strength analysis used to measure forelimb neuromuscular function showed no significant differences between the treated TKOc and CTRLs (Figure S2G). Finally, we did not detect significant differences in the lifespan of TKOc and CTRL mice (Figure S2H). Overall, these results show that our 5-day TAM treatment protocol induces a mild premature aging phenotype, maybe due to insufficient recombination efficiency. For this reason, we decided to perform an additional series of 5-day intraperitoneal injections of TAM at 5.5-months of age (Figure 2A).

**Figure 2.**
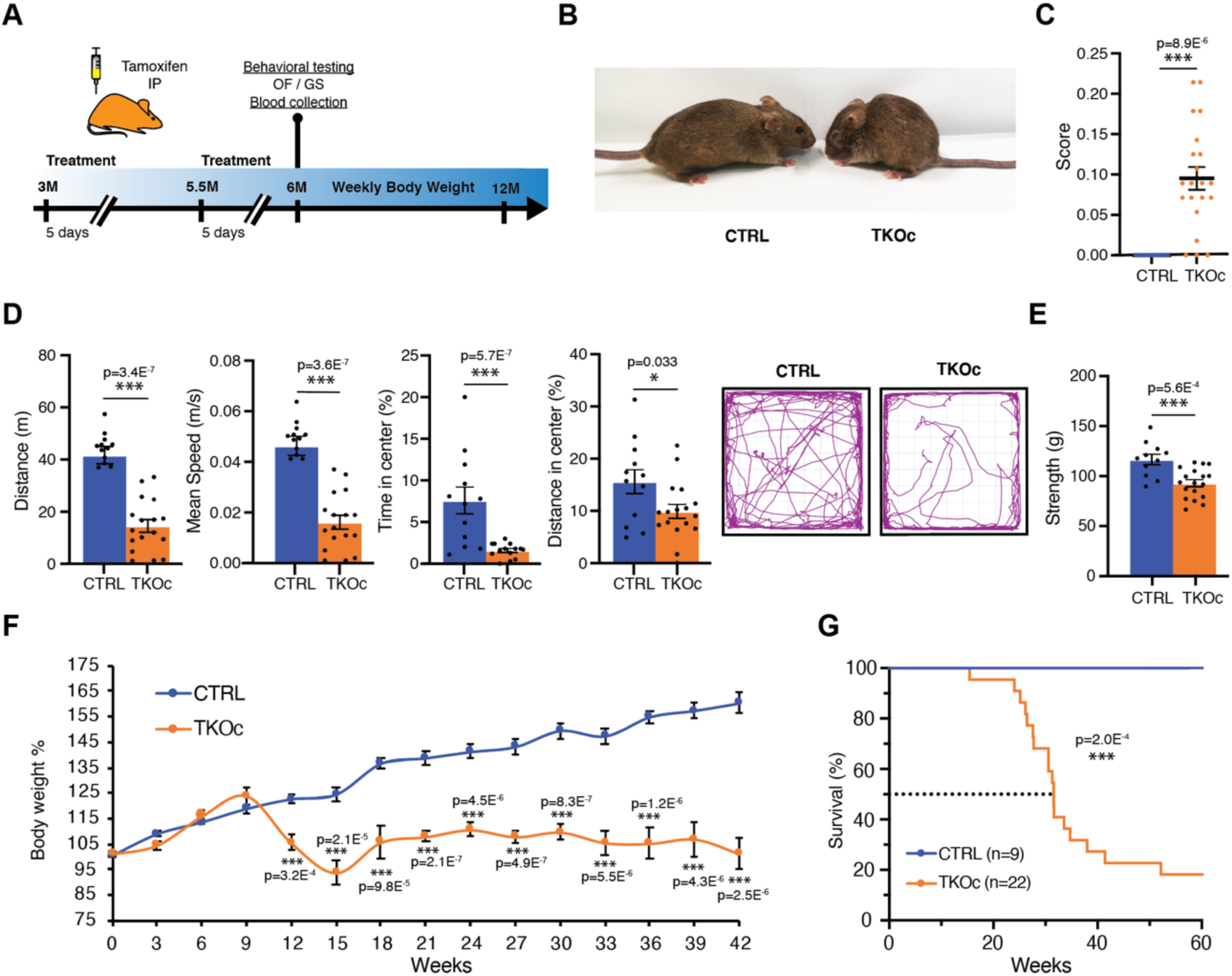
Loss of H3K9me3 *in vivo* leads to premature aging and decrease in lifespan. **(A)** Experimental design. **(B)** Image of CTRL and TKOc mice (6-month-old mice). **(C)** Frailty index of CTRL and TKOc mice (n = 13 to 21, 6-month-old mice). **(D)** Behavioral characterization of CTRL and TKOc mice (6 months). Open field exploration (15 minutes), quantification of distance travelled, mean speed, time spent and distance traveled in the center with a representative tracking trace (n = 12 to 14-17, 6-month-old mice). **(E)** Grip strength measured as average grip strength of three different trials CTRL and TKOc mice. (n = 11 to 17, 6-month-old mice). **(F)** Changes in body weight of CTRL and TKOc mice upon administration of tamoxifen **(G)** Kaplan-Meier survival curves for CTRL and TKOc mice (n = 9 to 22 mice, including males and females). Dots in all panels represent individual sample data. Survival curve data were analyzed by log-rank (Mantel-Cox test). Data are mean ± SEM. Statistical significance was assessed by Two-tailed Student’s t test. *p < 0.05; **p < 0.01; ***p < 0.001.

After the second round of TAM injections, we first assessed the percentage of recombination across various tissues. Our analyses revealed a lack of leakiness and a high recombination rate in proliferative tissues, including the skin, small intestine, and spleen. Additionally, we observed substantial recombination in skeletal muscle and brain tissues, with the lowest recombination rate in the liver (Figure S3A). Subsequently, we examined H3K9me3 protein levels in highly proliferative tissues of TKOc mice, specifically the skin and small intestine. Our findings indicated a significant reduction in H3K9me3 levels in TKOc mice compared to CTRLs (Figure S3B and S3C). Importantly, after the second round of TAM injections, TKOc mice displayed a severe aged appearance (Figure 2B). This premature aging phenotype was confirmed by a significant increase in FI scores in TKOc mice compared CTRL littermates (Figure 2C). Next, we evaluated activity by open field test and observed that TKOc mice had decreased activity and were slower than the CTRL mice (Figure 2D). Additionally, we noted that the TKOc mice spent substantially less time exploring the center of the arena, and traveled less, as compared to CTRL littermates (Figure 2D). Moreover, we found that treated TKOc mice had reduced grip strength compared to controls (Figure 2E). Combined with the appearance of frailty changes (noted in Figure 2C), a significant reduction in body weight was also observed in the treated TKOc mice, which was not recovered over time (Figure 2F). Most importantly, we observed a significant reduction in lifespan, with a media lifespan of only 30 weeks (Figure 2G). Altogether, our data demonstrates that loss of H3K9me3 in TKOc mice leads to premature aging.

### Degeneration of multiple organs is observed upon loss of H3K9me3

Given the observed premature aging phenotype of TKOc mice, we decided to conduct an in-depth characterization of the model at 6-8 months of age, a time point corresponding to the relative median survival of the TKOc strain. For this reason, we examined age-related features that are well characterized in the hematopoietic system, skin, small intestine and bone (Martin, Kirkwood and Potten, 1998; Ferguson *et al*., 2003; Russell-Goldman and Murphy, 2020). Interestingly, we observed changes in blood parameters including a notable reduction in the number of red blood cells (RBC), white blood cells (WBC) and the levels of hemoglobin (HGB), revealing an anemic condition compared to CTRL mice (Figure 3A; Figure S4A). In regard to the skin, compared with age-matched CTRLs, the skin of treated TKOc mice showed a pronounced loss of hypodermal fat and subepidermal thinning (Figure 3B). Furthermore, histological analysis of TKOc mice intestines revealed a reduction in the number and length of villi together with crypt deepening and ballooning (Figure 3C). Lastly, age-associated loss of bone thickness was observed in treated TKOc mice similar to physiological aging (Figure 3D). Bone microarchitecture analysis revealed that TKOc mice displayed a significant bone loss compared to littermate CTRLs, including decreased bone volume fraction (BV/TV), cortical thickness, cortical area and increase bone marrow area. Collectively, these findings strongly support that age-associated epigenetic alterations, such as the loss of H3K9me3, can drive aging in multiple organs.

**Figure 3.**
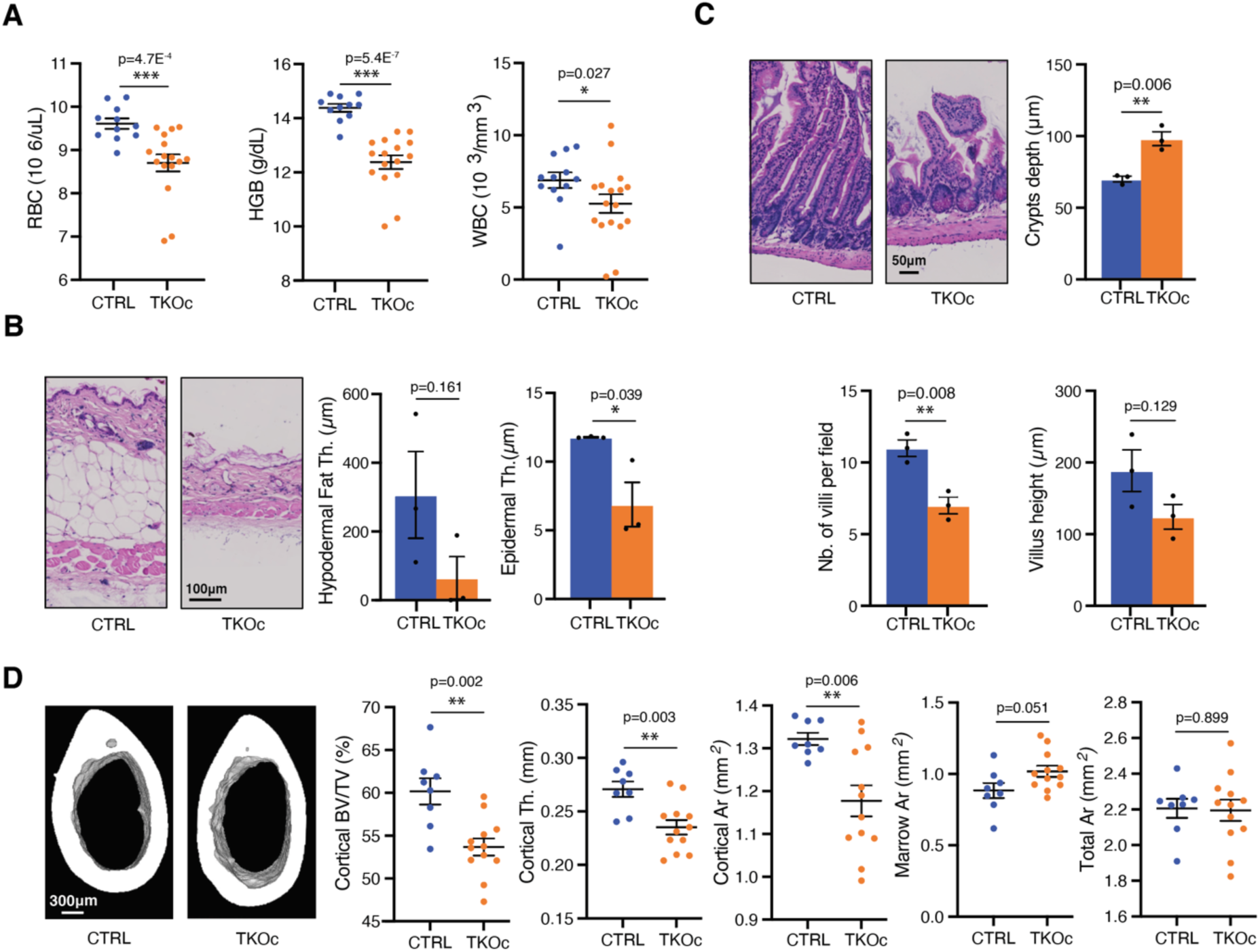
Age-associated organ degeneration results from H3K9me3 loss. **(A)** Hematological parameters in CTRL and TKOc mice (n = 11-12 to 16-17, 6-month-old mice). **(B)** Representative skin sections stained with hematoxylin and eosin (right) and quantification of hypodermal fat and epidermal thickness (left) in CTRL and TKOc mice. **(C)** Representative small intestine sections stained with hematoxylin and eosin (right) and quantification of crypt depth, number and height of the villus (left) in CTRL and TKOc mice. For the skin and small intestine at least 10 measurements were performed per animal. The graph shows mean values for n = 3 mice at 7-8 months of age. Scale bar, 100 or 50μm. **(D)** Representative micro-CT images (right) and quantitation of femur cortical bone (bottom; scale bar, 300μm) in CTRL and TKOc mice (n = 8 to12, 7-8-month-old mice). Data are mean ± SEM. Statistical significance was assessed by Two-tailed Student’s t test. *p < 0.05; **p < 0.01; ***p < 0.001.

### Loss of H3K9me3 leads to age-associated epigenetic and transcriptional changes

To further confirm that H3K9me3 loss results in premature aging at the molecular level, we used tissue-specific epigenetic clocks to assess DNA methylation age (DNAm) in TKOc and CTRL mice. Specifically, we analyzed both proliferative (skin, spleen, and small intestine) and post-mitotic tissues (skeletal muscle, liver, and brain) to account for potential differences in the levels of H3K9me3 recombination. Interestingly, accelerated DNAm age was detected in proliferative tissues of TKOc mice including the skin and small intestine as well as the spleen (Figure 4A). In addition, a tendency for accelerated aging was also observed in muscle, while no significant differences were detected in the liver and brain (Figure S5A). Moreover, the mean DNA methylation levels of TKOc mice was reduced in most proliferative tissues, most prominently in the small intestine and spleen, in line with the role of H3K9me3 in recruiting the DNA methylation maintenance machinery to the DNA replication fork during cell division. Loss of H3K9me3 therefore correlates with global DNA methylation loss and might contribute to the deregulation of related transcripts and de-repression of heterochromatin. (Figure S5B) Overall, our results indicate that the TKOc mouse model exhibits an accelerated epigenetic age and global loss of DNA methylation compared to age-matched controls.

**Figure 4.**
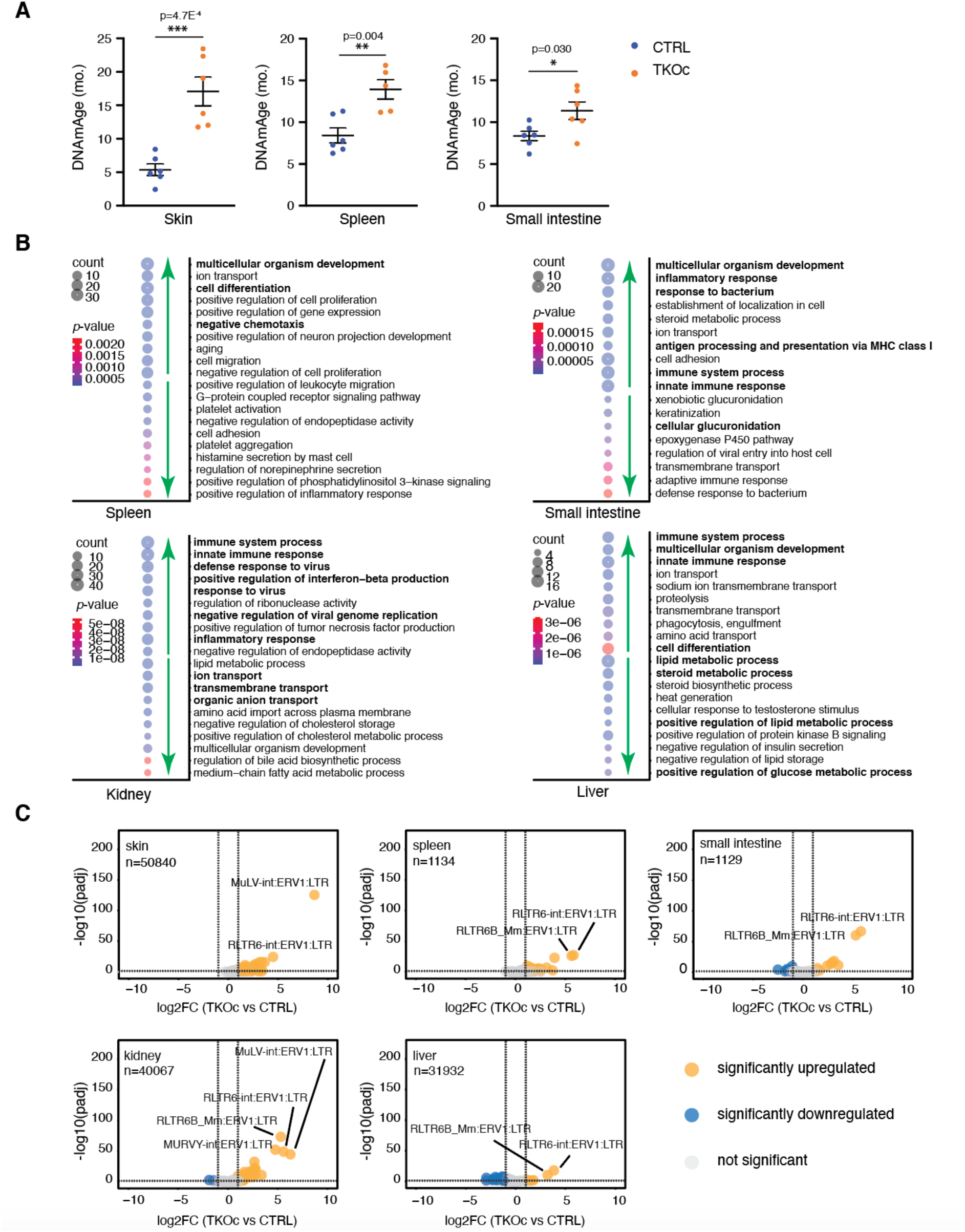
Accelerated epigenetic age and transcriptional dysregulation in TKOc mice. **(A)** Epigenetic age of skin, spleen and small intestine of mice TKOc mice compare their CTRL (n = 6 to 5-6, 7-8-month-old mice). Data are mean ± SEM. Statistical significance was assessed by Two-tailed Student’s t test *p < 0.05; **p < 0.01; ***p < 0.001. **(B)** Bubble plots of the top significant (up and down) GO terms in spleen, small intestine, kidney, and liver. **(C)** Volcano plots showing the TE transcripts that are significantly upregulated (in orange), downregulated (in blue) or unchanged (in gray) in skin, spleen, small intestine, kidney, and liver. Transcripts were determined to be significant if the adjusted *p-*value was < 0.05 and log_2_fold change was > 1. A few top upregulated transcripts are labeled in each tissue.

To perform a comprehensive analysis of transcriptomic changes resulting from the loss of H3K9me3, we generated total RNA-seq libraries from 7 different tissues from TKOc and CTRL mice and livers of 3-month-old young and 18-month-old aged C57BL6/JN mice (n=6 per group, equally distributed males and females, total n=96 libraries). The 7 different tissues were derived from both proliferative (spleen, small intestine, skin) and non-proliferative (liver, muscle, brain, and kidney) organs. Subsequently, we processed the raw FASTQ reads to enable both unique and multiple alignment with the mouse reference genome, which allowed us to generate counts from both genes and TEs.

Principal Component Analysis (PCA) of the transcriptomic data from TKOc and CTRL mice showed that the primary source of variation was the tissue type (Figure S5C). We then focused on the liver and performed a separate PCA analysis with TKOc, CTRL, young, and aged mice. Interestingly, the samples segregated by age on PC1 and by sex on PC2 with the TKOc samples showing a pro-aging transcriptome (Figure S5D, note shift to left similar to aged samples). Next, we used conventional DESeq2 analysis with uniquely mapped reads to identify differentially abundant mRNAs between TKOc and CTRL mice. For each tissue type, we then performed a Gene Ontology (GO) analysis (Huang, Sherman and Lempicki, 2009) to derive insight into functional pathways activated by H3K9me3 depletion (Figure 4B). Interestingly, several biological processes showed significant enrichments across tissues. These included immune/inflammatory response, response to virus, multicellular organism development, and cell differentiation. In contrast, downregulated mRNAs were related to tissues-specific function including glucuronidation in the small intestine, metabolic processes in the liver, and ion transport in the kidney. Notably, in our previous study of the aging liver from naturally aged mice, we observed similar GO terms up or downregulated (N. Yang *et al*., 2023).

Lastly, we used DESeq2 (Love, Huber and Anders, 2014) to identify differentially abundant TE transcripts between TKOc and CTRL tissues. Across the 7 different tissue types, there were more TEs detected in skin, muscle, liver, and kidneys than in spleen, small intestine, and brain (Figure 4C; Figure S5E, note n). More importantly, in all tissues derived from TKOc mice, there were more TEs that were significantly upregulated than downregulated (Figure 4C). We were curious to note that the top identified TEs in all tissues were ERVs (Figure 4C; Figure S5E, note labels). This observation prompted us to categorize all upregulated TEs to identify whether there was a preference for any specific family of TEs that are upregulated. Importantly, we found that among the top 50 most significantly upregulated TEs, ERVs were indeed the dominant family followed by LINEs showing the largest fold changes (Table S1 for details). This pattern of robust upregulation of TEs and the preference for ERVs was replicated in the livers of young and old C57BL/6JN mice (Figure S5E last panel; Table S1 for details). Overall, our comprehensive analyses of the transcriptome across both coding and non-coding regions of the genome highlight that H3K9me3 deregulation activates canonical age-related transcriptional pathways including upregulation of TEs. Thus, demonstrating that TKOc mice show features of premature aging at the molecular and cellular level.

## DISCUSSION

The molecular mechanisms and causal relationships underlying age-related transcriptional changes and the loss of heterochromatin during aging has largely remained elusive. According to the “’heterochromatin loss” theory of aging, the degradation of constitutive heterochromatin at the nuclear periphery, marked by H3K9me3, is believed to play a pivotal role during the aging processes driving pro-aging cellular phenotypes (Kane and Sinclair, 2019; N. Yang *et al*., 2023). Previous investigations have documented the age-associated decline of H3K9me3 in multiple models including *D. melanogaster*, *C. elegans,* as well as human cells from old individuals and premature aging patients (Scaffidi and Misteli, 2006; Larson *et al*., 2012; Ni *et al*., 2012; Zhang *et al*., 2015a). Consequently, the age-related reduction in H3K9me3 has been postulated as a key contributing factor to the aging process.

H3K9me3 constitutes a post-translational modification recognized for its involvement in regulating diverse biological processes, particularly in the establishment of transcriptionally silent heterochromatin. Proposed models align with the idea that H3K9me3 loss accentuates the aging process through the dysregulation of chromatin organization. Through the simultaneous depletion of Setdb1 and Suv39h1/2, methyltransferases crucial to the formation of constitutive heterochromatin, our model analyzes consequential transcription changes including a potential source of genomic instability by the activation of endogenous mobile genetic elements, specifically transposable elements (He *et al*., 2019; Gorbunova *et al*., 2021).

In this study, we aim to demonstrate the causative role of H3K9me3 as driver of aging. Towards this goal, we circumvent the detrimental consequences of the constitutive loss of H3K9me3 during the embryonic development by generating a tamoxifen inducible mouse model that enables the depletion of H3K9me3 in adulthood (Nicetto *et al*., 2019a). Our findings demonstrate that the induced loss of H3K9me3 during adulthood leads to the manifestation of premature aging phenotypes, characterized by an increased frailty, accelerated aging across diverse tissues, and a reduction in lifespan. At the molecular level, loss of H3K9me3 results in de-repression of TEs, upregulation of lineage-inappropriate developmental genes, and downregulation of mRNAs specifying tissue function. Through the direct targeting of H3K9me3, we provide substantiating evidence that loss of epigenetic information drives mammalian aging. Importantly, in alignment with the interconnectivity between different hallmarks of aging, DNA damage globally impacts chromatin structure. In this line, a recent study demonstrated that the repair of DNA breaks can lead to epigenetic erosion and a global reduction in chromatin compaction, ultimately resulting in premature aging (J.-H. Yang *et al*., 2023). Similarly, we have recently shown that DNA-repair deficient premature aging models display accelerated epigenetic age (Perez *et al*., 2024).

While the decline in H3K9me3 is evident across various species during aging, this pattern is also contingent on the specific tissues and cell types (Ocampo *et al*., 2016; Snigdha *et al*., 2016; Rodríguez-Matellán *et al*., 2020). Consistent with this observation, we observed a more pronounced increase in biological age, particularly in proliferative tissues: spleen, skin, and small intestine, which might undergo a faster depletion of H3K9me3 by the lack of methyltransferases that can restore this mark after cell division. However, irrespective of proliferative potential, we found that loss of H3K9me3 increases the expression of TEs, particularly ERVs and LINEs, in all 7 tissues examined from TKOc mice. Interestingly, ERVs have recently been shown to be upregulated in aged murine, primate and human cells and organs, as well as serum from older individuals (Liu *et al*., 2023). Moreover, ERVs have been recently to lead to cellular senescence and tissue aging (Liu *et al*., 2023). Similarly, several reports have demonstrated that LINE-1 elements can drive aging in mice (De Cecco *et al*., 2019; Simon *et al*., 2019). These results were also recapitulated when analyzing RNA-seq data from aged mice livers. Importantly, LINE-1 and ERVs both result in activation of the innate immune system and consequently the release of senescence-associated secretory phenotype (SASP) factors. Indeed, our conventional analysis of differential mRNAs from annotated genes show a strong innate immune activation signature. Conversely, downregulated mRNAs were mostly related to the tissue type indicating a loss of tissue function.

Lastly, we hypothesize that interventions known to slow the process of aging process leading to healthy lifespan might mitigate associated hallmarks of aging including epigenetic dysregulation. In this line, restoration of H3K9me3 levels is an early event observed during the rejuvenation of age-associated phenotypes by cellular reprogramming, and preventing H3K9me3 restoration using an H3K9 methyltransferase inhibitor is sufficient to block the amelioration of additional age-associated phenotypes (Ocampo *et al*., 2016). Similarly, therapeutic interventions that promote longevity have been shown to reduce the expression of repetitive elements (RE) (Wahl *et al*., 2021).

In conclusion, our data strongly support the notion that changes in the epigenome, including the age-associated loss of H3K9me3, can drive the aging processes in mammals, and therefore reinforce the role of epigenetic dysregulation as an important hallmark of aging. The TKOc mice generated in this study could potentially serve as a valuable experimental model for in-depth investigation of the molecular mechanisms of aging. Lastly, the loss of H3K9me3 might represent a novel target for the potential development of therapeutic interventions aiming at the prevention of age-associated diseases and extension of healthy lifespan.

## ACKNOWLEDGMENTS

The authors thank all members of the Ocampo and Payel Sen laboratory for helpful discussions. We are very grateful to Kenneth Zaret for kindly providing use the TKO mice. We thank the teams of mouse facilities at the University of Lausanne including Francis Derouet (head of the animal facility at Epalinges) and L. Lecomte (head of the animal facility of the Department of Biomedical Sciences). We thank the Mouse Pathology Facility of the University of Lausanne for tissue processing and stainings. We acknowledge the funding from NIA intramural program and the computational resources of the NIH HPC Biowulf cluster (http://hpc.nih.gov).

## AUTHOR CONTRIBUTIONS

C.M., A.P., A.O. conceived the project. C.M., N.Y., G.D.-M., A.A.C, O.N. performed experiments. C.B. and M.C.M. performed genotyping. A.V.A., M.C.M., C.Y.M., S.P. performed sample collection. N.Y. and Y.P. performed RNA extraction from tissues. L.S. guided for immunofluorescence imaging. N.Y., P.S., S.L., F.V.M. did the bioinformatics analysis. A.M., R.B., S.H. did the epigenetic clock analysis. P.S. and A.O. contributed to personnel supervision. C.M., N.Y., P.S., A.O. wrote the original draft. C.M., N.Y., P.S., A.O., G.D.-M., A.V.A., S.P wrote, review and edit from the original draft.

## DATA AVAILABILITY

The authors confirm that data supporting the findings of this study are available within the article and its Supplementary Information, or are available from the corresponding author upon reasonable request. Materials: The mouse model described in this work will be made available to investigators through an institutional or third-party Material Transfer Agreement (MTA) upon reasonable request. Data: Raw sequencing reads related to tissue analysis were deposited in the Gene Expression Omnibus (GEO) under the project number (GSE262109).

## METHODS

### Animal housing

All the experimental procedures were performed in accordance with Swiss legislation after the approval from the local authorities (Cantonal veterinary office, Canton de Vaud, Switzerland). Animals were housed in groups of five mice per cage with a 12hr light/dark cycle between 06:00 and 18:00 in a temperature-controlled environment at 25°C and humidity between 40 and 70% (55% in average), with free access to water and food. Transgenic mouse models used in this project were generated by breeding and maintained at the Animal Facility of Epalinges and the Animal Facility of the Department of Biomedical Science of the University of Lausanne.

### Mouse strains

The TKOCAGCre mouse strains were generated by breeding the TKO strain, triple conditional knockout for the three H3K9me3 methyltransferases (Suv39h2, Suv39h1, Setdb1), previously generated by Professor Kenneth Zaret (Nicetto *et al*., 2019b) with CAG-CreER^TM^ mice Stock No 004682. The final Homo-TKOCAGCre mouse strain is a quadruple transgenic conditional knockout mouse carrying: Setdb1 Flox/Flox: Loxp sites flanking exon 15-16 of Setdb1 gene (Chr.3). Suv39h1 Flox/Flox: Loxp sites flanking exon 3-5 of Suv39h1 gene (Chr.X). Suv39h2 KO/KO: deletion in the SUV39H2 gene (Chr. 10). CAG-CreER Cre/+: Insertion of CAG-CreER (Chr.3), for the tamoxifen-inducible CRE-mediated recombination system (Loxp sites). TKOCAGCre littermates not expressing Cre were used as control mice along the study.

### Tamoxifen administration

Tamoxifen (Sigma, T5648) of TKOCAGCre mice was performed at 3 months of age and repeated at 5.5 months. Tamoxifen was administrated intraperitoneally at 67mg/kg for 5 consecutive days.

### Mouse monitoring and euthanasia

All mice were monitored at least three times per week. Upon Tamoxifen injection, mice were monitored twice a week to evaluate their activity, posture, alertness, body weight and presence of tumors or wound. Mice were euthanized according to the criteria established in the scoresheet. We defined lack of movement and alertness, presence of visible tumors larger than 1cm^3^ or opened wounds and body weight loss of over 30% as imminent death points. Both genders were used for survival, body weight experiments, tissue and organ collection. Animals were sacrificed by CO_2_ inhalation (6 min, flow rate 20% volume/min). Subsequently, before perfusing the mice with saline, blood was collected from the heart. Finally, multiple organs and tissues were collected in liquid nitrogen and used for DNA, RNA, and protein extraction, or placed in 4% formalin for histological analysis.

### Behavior

Behavioral characterization was performed on both males and females, at the age of 6 and 12 months. Open field (OF:) Locomotor activity and anxiety-like behavior of adult mice were evaluated in an open field arena. Briefly, mice were individually placed in the center of a Plexiglas boxes (sides 45 cm, height 40 cm, Harvard Apparatus, 76-0439). Mice movements were recorded for 15 minutes. Recording was done with a USB camera (Stoelting Europe, 60516) and then analyzed using ANY-maze video tracking software (ANY-maze V7.11, Stoeling). Grip strength test (GS): to measure muscular strength, a mouse was held by the tail and allowed to grip a mesh grip with the front paws (Harvard Apparatus, 76-1068). Three measurements minimum per trial were performed for each animal, with a few seconds resting period between measurements.

### Frailty Index assessment

The Frailty Index (FI) was adapted from the previously described score (Whitehead *et al*., 2014). For each mouse 28 health-related deficits were assessed going across the integument, physical/musculoskeletal, ocular/nasal, digestive/urogenital and respiratory systems were scored as 0, 0.5 and 1 based on the severity of the deficit. Total score across the items was divided by the number of items measured to give a frailty index score between 0 and 1.

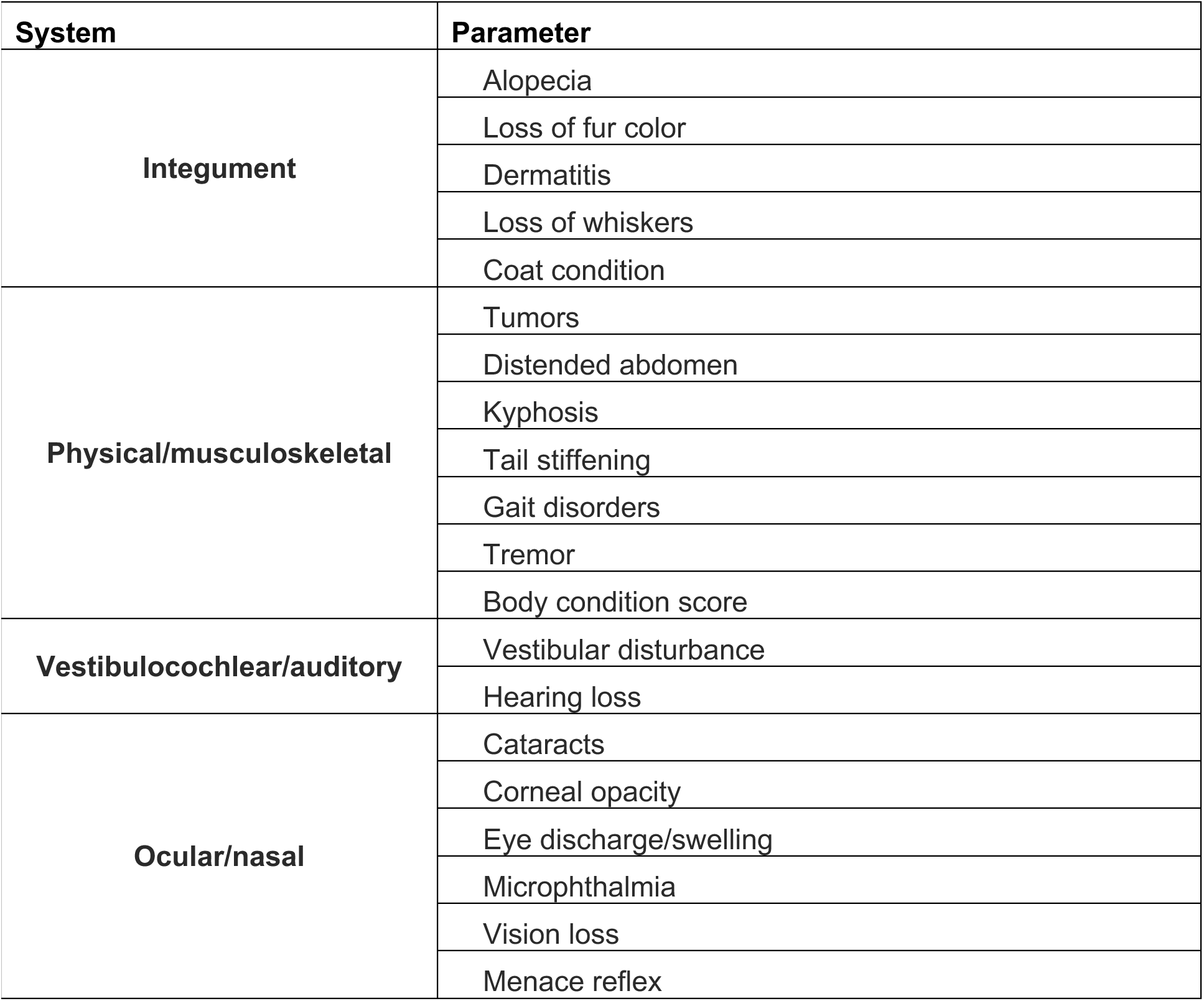
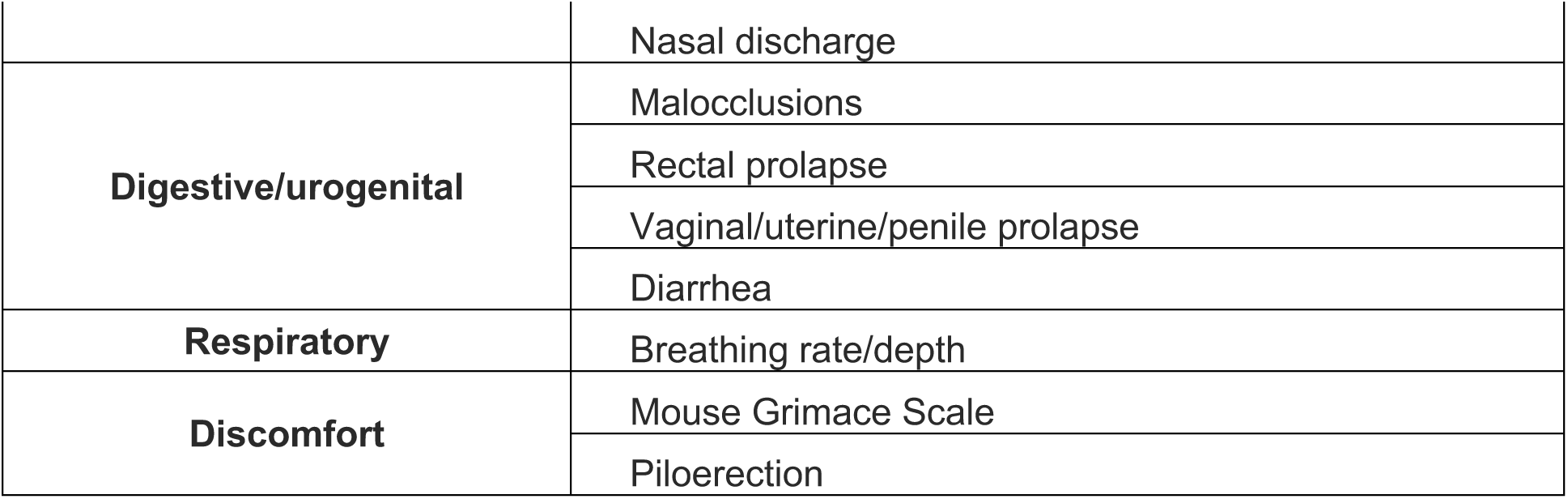

| System | Parameter |
| --- | --- |
| Integument | Alopecia |
|  | Loss of fur color |
|  | Dermatitis |
|  | Loss of whiskers |
|  | Coat condition |
| Physical/musculoskeletal | Tumors |
|  | Distended abdomen |
|  | Kyphosis |
|  | Tail stiffening |
|  | Gait disorders |
|  | Tremor |
|  | Body condition score |
| Vestibulocochlear/auditory | Vestibular disturbance |
|  | Hearing loss |
| Ocular/nasal | Cataracts |
|  | Corneal opacity |
|  | Eye discharge/swelling |
|  | Microphthalmia |
|  | Vision loss |
|  | Menace reflex |
|  | Nasal discharge |
| **Digestive/urogenital** | Malocclusions |
|  | Rectal prolapse |
|  | Vaginal/uterine/penile prolapse |
|  | Diarrhea |
| **Respiratory** | Breathing rate/depth |
| **Discomfort** | Mouse Grimace Scale |
|  | Piloerection |

### Hematological analysis

Blood was collected from the temporal vein in potassium EDTA microtrainer tubes. Complete blood count was performed in Heska Element HT5 hematology analyzer.

### Bones analysis

Bone microarchitecture was evaluate using a SkyScanner 1276 (Bruker, Belgium). 0.25 mm Al filter was used with a voltage of 200 kV and a current of 55 mA. To avoid drying samples were wrapped in paper towels soaked in PBS and scanned inside a drinking straw sealed on both ends. Voxel size for both applications was set at 10×10×10 µm3.

Bone microarchitecture was evaluated according to the ASBMR guidelines (Bouxsein *et al*., 2010) using a custom CTan (Bruker, Belgium) script for automatic segmentation of trabecular bone in the distal femoral VOI, which was set 100 slices proximal to the distal growth plate and extended 200 slices towards the femoral diaphysis (slice thickness of 0.010 mm). The threshold used to binarize the calcified tissue was 40 on a 0-255 scale. Reconstruction of the scans was performed using NRecon (Bruker, Belgium) and further analysis were performed using CTan (Bruker, Belgium) with the minimum for CS to image conversion set at 0 and maximum set at 0.14. For the analysis of cortical parameters, the midpoint of the femur was determined, and the VOI was defined as the bone 50 slices (slice thickness of 0.010 mm) distal and proximal of the slice corresponding to the midpoint of the bone. All other parameters were kept the same as for the analysis of trabecular bone.

### DNA extractions

Total DNA was extracted from tissues using Monarch Genomic DNA Purification Kit (New England Biolab, T3010L). Tissues were cut into small pieces to ensure rapid lysis. Total DNA concentrations were determined using the Qubit DNA BR Assay Kit (Thermofisher, Q10211).

### DNA methylation clock

Analysis of epigenetic age was done in collaboration with the Clock Foundation. The mouse clock was developed in ref. (Mozhui *et al*., 2022). To analyze the epigenetic age, we used for skin, spleen, small intestine, skeletal muscle, liver and brain the following mouse clocks: “UniversalClock3Skin”, “UniversalClock2Blood”, “UniversalClock2Pan”, “DNAmAgeMuscleFinal”, “DNAmAgeLiverFinal”, “DNAmAgeCortexFinal”. The mouse methylation data were generated on the small and the extended version of HorvathMammalMethylChip (Arneson *et al*., 2022). We used the SeSaMe normalization method (Zhou *et al*., 2018). We used the noob normalization method implemented in the R function preprocessNoob.

### Overall DNA methylation

Analysis was done by first removing probes with only background signal in a high proportion of samples. This was done by retaining only those probes that have a detection p-value of 0.05 or greater in more than 90% of the samples. Afterwards, we eliminated probes located on the X and Y chromosomes. Subsequently, for each sample, we calculated the median of the beta-values from the remaining probes to estimate the overall DNA methylation level.

### Immunohistochemistry

Mice were euthanized with CO2 and multiple tissues and organs were collected, placed in 4% formalin (Sigma, 252549) overnight, and then immersed in 30% sucrose in phosphate buffered saline (PBS) for 72 h. Subsequently, samples were paraffin-embedded with a Leica ASP300S tissue processor (Leica, Heerbrug, Switzerland), sections prepared with a Microm HM 335 E microtome (Thermo Scientific,Walldorf, Germany) and mounted on Superfrost Plus slides (Thermo Scientific). Next, slides were deparaffinized and rehydrated with xylol and alcohol. Each section was routinely stained with hematoxylin and eosin, mounted on glass slides, and examined. For the analysis of H & E staining, the epidermal, subepidermal thickness, Crypt depth, Villus height, Number Villus were quantified in the skin and the small intestine in ten different regions per animal and means of the ten regions were calculated. For H3K9me3 immunostaining, the intensity was quantified in the small intestine and skin in four different regions per animal. Antibody used was rabbit Cell Signaling: Tri-Methyl-Histone H3 (Lys9) (D4W1U).

### RNA extraction

For cells total RNA extracted using Monarch Total RNA Miniprep Kit (New England Biolab, T2010s). Total DNA concentrations were determined using the Qubit RNA BR Assay Kit (Thermofisher, Q10210).

### cDNA synthesis

cDNA synthesis was performed by adding 4 μL of iScript™ gDNA Clear cDNA Synthesis (Biorad, 1725035BUN) to 500ng of RNA sample and run in a Thermocycler (Biorad, 1861086) with the following protocol: 5 min at 25°C for priming, 20 min at 46°C for reverse transcription, and 1 min at 95°C for enzyme inactivation.

### Semiquantitative RT-PCR

DNA was amplified using DreamTaq Green PCR Master Mix 2X (Thermofisher, K1081) following the amplification protocol: 3 min at 95°C + 33 cycles (30 s at 95°C + 30 s at 56 or 60 °C + 1 min at 72°C) + 5 min at 72°C. PCR products were loaded and run in an agarose (1.6%) gel containing ethidium bromide (Carlroth, 2218.1). Images were scanned with a gel imaging system (Genetic, FastGene FAS-DIGI PRO, GP-07LED). Setdb1 and Suv39h1 recombination were detected using the following primers: Setdb1forward: 5’- CAGCTTGGAGGAATTGGTTC-3’ Setdb1 reverse 1: 5’- TTTCTTTGCCTTTGAGATGGA-3’ Setdb1 reverse 2: 5’- TACCATACCCACTAACACTTTGC-3’, Suv39h1 forward: 5’-GGAGCCCACTGAAAGTAGCA-3’, Suv39h1 reverse 1: 5’- ACTCCAGCCCCTCCTTTTT-3’ Suv39h1 reverse 2: 5’- GGTCAGGCTAGAAAACACAAGG-3’.

### qRT-PCR

cDNA was diluted 1:5 using nuclease free water and stored at - 20°C. qRT-PCR was performed in a Quantstudio 12K Flex Real-time PCR System instrument (Thermofisher) using SsoAdvanced SYBR Green Supermix (Bio-Rad, 1725274) in a 384-well PCR plate (Thermofisher, AB1384). Forward and reverse primers were used at a ratio 1:1 and final concentration of 5 µM with 1ul of cDNA.

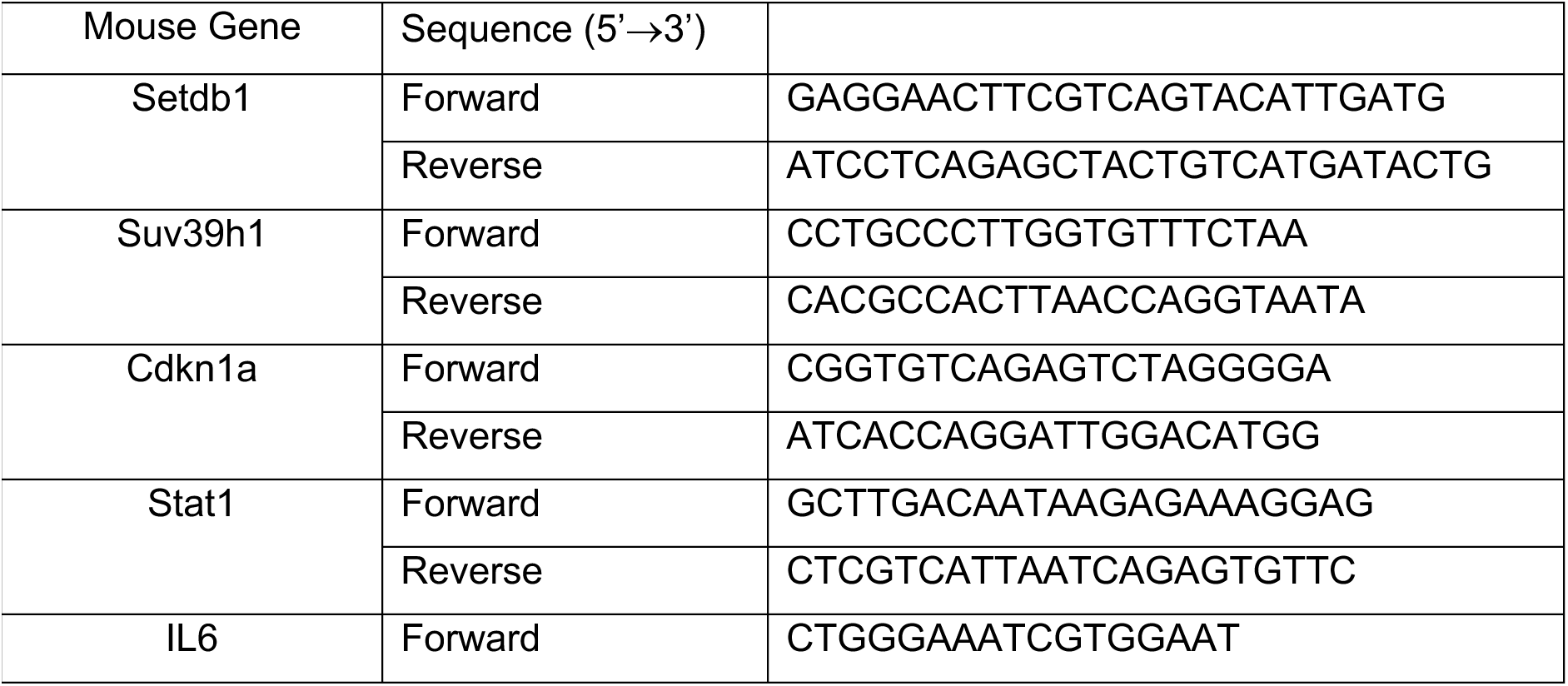
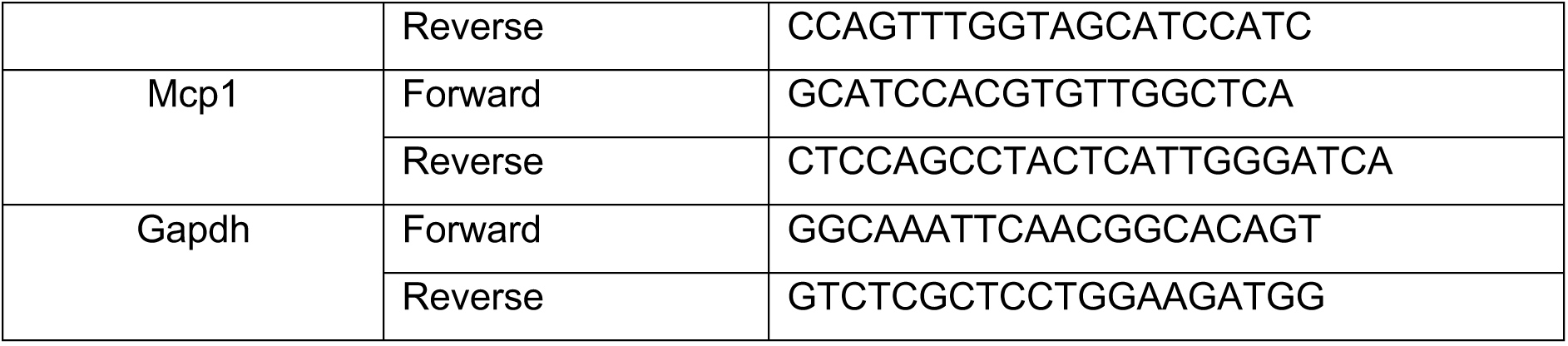

### Cell culture

Mouse tail-tip fibroblasts (TTFs) were freshly extracted using Collagenase I (Sigma, C0130) and Dispase II (Sigma, D4693) and cultured in DMEM (Gibco, 11960085) containing non-essential amino acids (Gibco, 11140035), GlutaMax (Gibco, 35050061), Sodium Pyruvate (Gibco, 11360039) and 10% fetal bovine serum (FBS, Hyclone, SH30088.03) at 37°C in hypoxic conditions (3% O2). Subsequently, fibroblasts were passaged and cultured according to standard protocols. For activation of the Cre recombination TTFs were treated with 0.1 μM 4-Hydroxytamoxifen (4-OHT) for 6 days and subsequently cultured in medium was switched to one without 4-OHT.

### Senescence-associated β-galactosidase assay

The senescence-associated beta-galactosidase (SA-βgal) assay was carried out following the method outlined by Debacq-Chainiaux et al. (2009). In summary, cells plated on glass coverslips underwent light fixation using a 3% paraformaldehyde and 0.2% glutaraldehyde solution in PBS buffer for 5 minutes. After removal of the fixation solution, the wells were washed multiple times and stained overnight at 37°C in a CO2-free incubator. The staining solution consisted of 40 mM citric acid/Na phosphate buffer, 5 mM K4[Fe(CN)6]·3H2O, 5 mM K3[Fe(CN)6], 150 mM sodium chloride, 2 mM magnesium chloride, and 1 mg/mL X-gal (Roth, 2315.1), with a pH range of 5.9-6.0. Subsequently, the coverslips were stained with DAPI and subjected to a standard immunofluorescence protocol. Bright-field microscopy was used to capture images, and the proportion of β-Gal-positive cells was quantified.

### Immunofluorescence staining

Cells were washed with fresh PBS and then fixed with 4% paraformaldehyde (Roth, 0964.1) in PBS at room temperature (RT) for 15 minutes. After fixation, cells were washed 3 times, followed by a blocking and permeabilization step in 1% bovine serum albumin (Sigma, A9647-50G) in PBST (0.2% Triton X-100 in PBS) for 60 min (Roth, 3051.3). Cells were then incubated at 4°C overnight with appropriate primary antibody, washed in PBS, followed by secondary antibody incubation with DAPI staining at RT for 60 min. Coverslips were mounted using Fluoromount-G (Thermofisher, 00-4958-02), dried at RT in the dark for several hours, stored at 4°C until ready to image and −20°C for long-term.

### Immunofluorescence imaging

Confocal image was acquired using the Ti2 Yokogawa CSU-W1 Spinning Disk (Nikon), using the 100X objective and with 15 z-sections of 0.3 μm intervals. The following lasers were used (405 nm and 488 nm) with a typical laser intensity set to 5-10% transmission of the maximum intensity for H3K9me3. Exposure time and binning were established separately to assure avoidance of signal saturation.

### Antibodies and compounds

Antibodies were provided from the following companies. Cell Signaling: Tri-Methyl-Histone H3 (Lys9) (D4W1U), Sigma: anti-β-Actin (A2228); Thermofisher: anti-Rabbit (A32790); Agilent: anti-Rabbit Immunoglobulins/HRP (P0448), anti-Mouse Immunoglobulins/HRP (P0447); Roth: DAPI (6843.1)

### Western blot

Cell and tissue were lysed in RIPA buffer (50 mM Tris pH 7.5, 0.5 mM EDTA, 150 mM NaCl, 1% NP40, 0.1% SDS), protease inhibitors and ceramic beads using a MagNa lyser instrument (Roche). To the lysate 10% SDS was added to bring up the SDS concentration to 1%. The homogenate was sonicated for 10 minutes with Bioruptor, 30s on and 30s off then centrifuged at 21,000 g for 10 min at 4°C. The resulting supernatants were collected, and protein content determined by Quick Start Bradford kit assay (Bio-Rad, 500-0203). 5-15 μg of total protein was electrophoresed on 10% SDS– polyacrylamide gel, transferred to a nitrocellulose blotting membrane (Amersham Protran 0.45 μm, GE Healthcare Life Sciences, 10600002) and blocked in TBS-T (150 mM NaCl, 20 mM Tris–HCl, pH 7.5, 0.1% Tween 20) supplemented with 5% Bovine Serum Albumin (BSA). Membranes were incubated overnight at 4°C with the H3K9me3 primary antibody in TBS-T supplemented with 5% BSA, washed with TBS-T and next incubated with secondary HRP-conjugated anti-rabbit IgG (1:2,000, DAKO, P0448) for 1 hour at room temperature and developed using the ECL detection kit (Perkin Elmer, NEL105001EA).Antibodies: mouse OCT-3/4 (C-10) (1:3000, Cell Signaling, 13969); mouse β-Actin (AC-74) (1:10,000, Sigma-Aldrich, A2228).

### RNA-sequencing

Total RNA was extracted by using the RNeasy Fibrous Tissue Mini Kit (Qiagen) with DNase treatment from 7 different tissues (liver, muscle, skin, spleen, small intestine, brain, and kidney) from TKOc and CTRL mice and livers of 3-month-old young and 18-month-old aged C57BL6/JN mice (n=6 per group, equally distributed males and females, total n=96 libraries) using a QIACube (Qiagen). Briefly, frozen tissue samples were transferred to 2 mL tubes (Sarstedt) containing 480 µL of RLT (RNA lysis buffer) (Qiagen) plus dithiothreitol (DTT, Teknova) buffer from the Qiagen kit, and 1 mm diameter Zirconia beads (Biospec) were added to make a total volume of 800 µL. The tube was processed using the Precellys 24 homogenizer, followed by centrifugation at 17,000 RCF (g) for 5 min at 4°C. The resulting homogenate was transferred to 1.5 mL Eppendorf tubes for Proteinase K treatment, and subsequently used for RNA isolation. A DNase digestion step was incorporated to remove contaminant genomic DNA. The quality and quantity of RNA were assessed using the Tapestation and the RNA Screen Tape (Agilent). RNA integrity numbers (RINs) ranged between 2.2-9.1 with the spleen having the lowest RINs due to high amounts of RNase.

∼250 ng of total RNA from each sample was used as input to prepare libraries for total RNA-seq with Zymo-Seq RiboFree Total RNA Library Kit (Zymo Research) following the manufacturer’s protocol. The quality and quantity of the libraries were assessed using the High Sensitivity DNA 1000 Screen Tape (Agilent) on a Tapestation (Agilent). The RNA-seq libraries were pooled and paired-end sequenced on the NovaSeq 6000 platform (Illumina) using an S2 200 cycle kit (100 paired-end). We obtained ∼45 million paired-end reads per sample.

### RNA-seq analysis

For conventional gene-based analysis, Illumina sequencing reads (∼45 million paired-end reads per sample) were de-multiplexed using bcl2fastq/2.20.0.422. Reads were trimmed to remove adapter sequences using trimmomatic/0.39 (https://www.bioinformatics.babraham.ac.uk/projects/trim_galore/). The quality of the resulting FASTQs was assessed using FASTQC/0.11.9 (https://www.bioinformatics.babraham.ac.uk/projects/fastqc/) and reads were aligned to the mouse reference genome (assembly GRCm38/mm10) using STAR/2.7.10b (PMID: 23104886). BAM files were sorted and indexed using samtools/1.17 (PMID: 19505943) and duplicates were removed using picard/3.1.0 (https://broadinstitute.github.io/picard/). The BAM files were then filtered to retain alignments with a minimum mapping quality of 10 using samtools/1.17 (PMID: 19505943). The featureCounts function of the Rsubread R package/2.16.0 was used to estimate counts for all transcripts. Differential gene expression analysis was performed using the R Bioconductor package, DESeq2/1.42.0.

For transposable element (TE) expression, trimmed FASTQs were aligned allowing for multi-mapping using STAR/2.7.10b (PMID: 23104886) with unsorted BAM files as output. This was followed by assessment of significantly altered transcripts with the TE transcripts function in TEToolkit/2.2.3 (PMID: 29508296).

### Gene Ontology (GO) analysis and bubble plots

GO analysis was performed using DAVID/2021 (PMID: 19131956) with expressed genes as background for each tissue. The top 10 significant GO terms for upregulated and downregulated mRNAs for the Biological Process category are reported. The bubble sizes are scaled on normalized counts and colored on *p*-value. For bubble plots showing TE expression, the bubble sizes are scaled on log_2_fold change, colored by category, and with −log_10_ *p-*values on the x-axis.

### Volcano plots

Volcano plots of TE expression changes were plotted using the VolcaNoseR online tool (PMID: 33239692).

### PCA plots

PCA plot was generated in R/4.0 with the TEtranscripts output using DESeq2 and then plotted with ggplot2.

### Statistical analysis

Statistical analyses were conducted using GraphPad Prism (version 10). Outliers were identified and excluded using the ROUT method (5% threshold). For comparisons between two groups, unpaired two-tailed Student’s t-tests were used for data that met normal distribution criteria, and the Mann-Whitney U test was used for data that did not meet normal distribution criteria, as determined by the Shapiro–Wilk normality test. Data are presented as mean ± SEM (standard error of the mean). For comparisons involving more than three groups, One-Way ANOVA was used for data that met normal distribution criteria, and the Kruskal-Wallis test was used for data that did not meet normal distribution criteria, as determined by the Shapiro–Wilk normality test. Data are presented as median with interquartile range. Unless otherwise specified, ‘n’ represents the number of individual biological replicates, and each sample is represented as a dot on the graphs.

## Supplemental Information

### Supplementary Figure Titles and Legends

**Figure S1:**
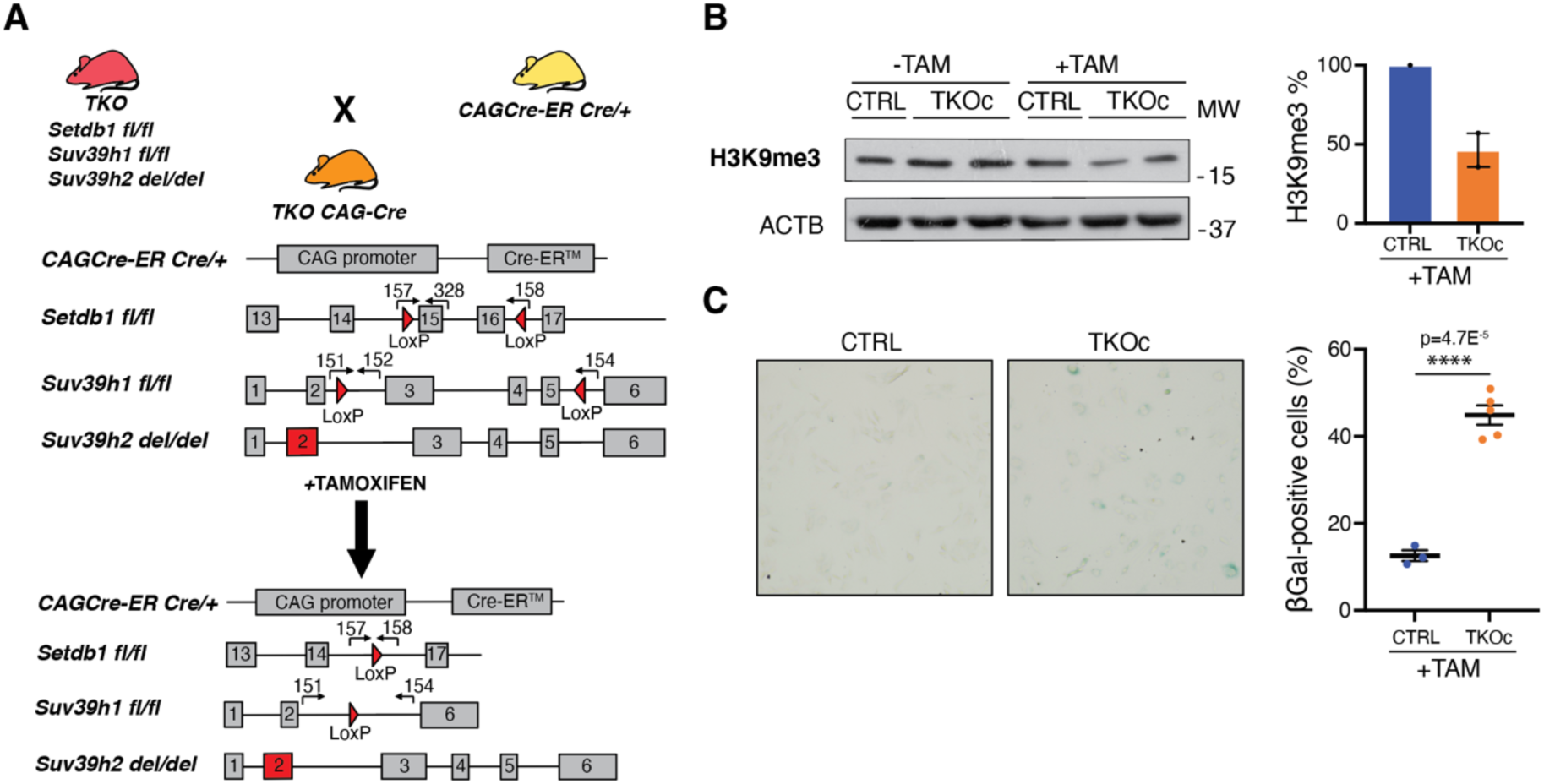
Genetic strategy for the generation of TKOc mice and loss of H3K9me3 *in-vitro*. **(A)** Detailed schematic representation of the genetic strategy to generate the quadruple transgenic TKOCAGCre mouse strain carrying the insertion of CAG-CreER (Chr.3), for the tamoxifen-inducible CRE-mediated recombination system (Loxp sites), Setdb1 gene insertion of LoxP sites in the intron 14 and 16 (Chr.3), Suv39h1 gene insertion of LoxP sites in the intron 2 and 5 (Chr.X), Suv39h2 gene deletion at Exon2 (Chr. 10). **(B)** Western blot of H3K9me3 in CTRL and TKOc tail-tip fibroblasts with or without TAM treatment. H3K9me3 quantification in CTRL and TKOc after treatment. **(C)** Senescence-associated β-galactosidase (SA-β-gal) staining in CTRL and TKOc tail-tip fibroblasts with TAM treatment. Quantification of SA-β-gal positive cells for CTRL and TKOc after treatment. Data are mean ± SEM. Statistical significance was assessed by Two-tailed Student’s t test *p < 0.05; **p < 0.01; ***p < 0.001 of 3 or 5 replicates.

**Figure S2:**
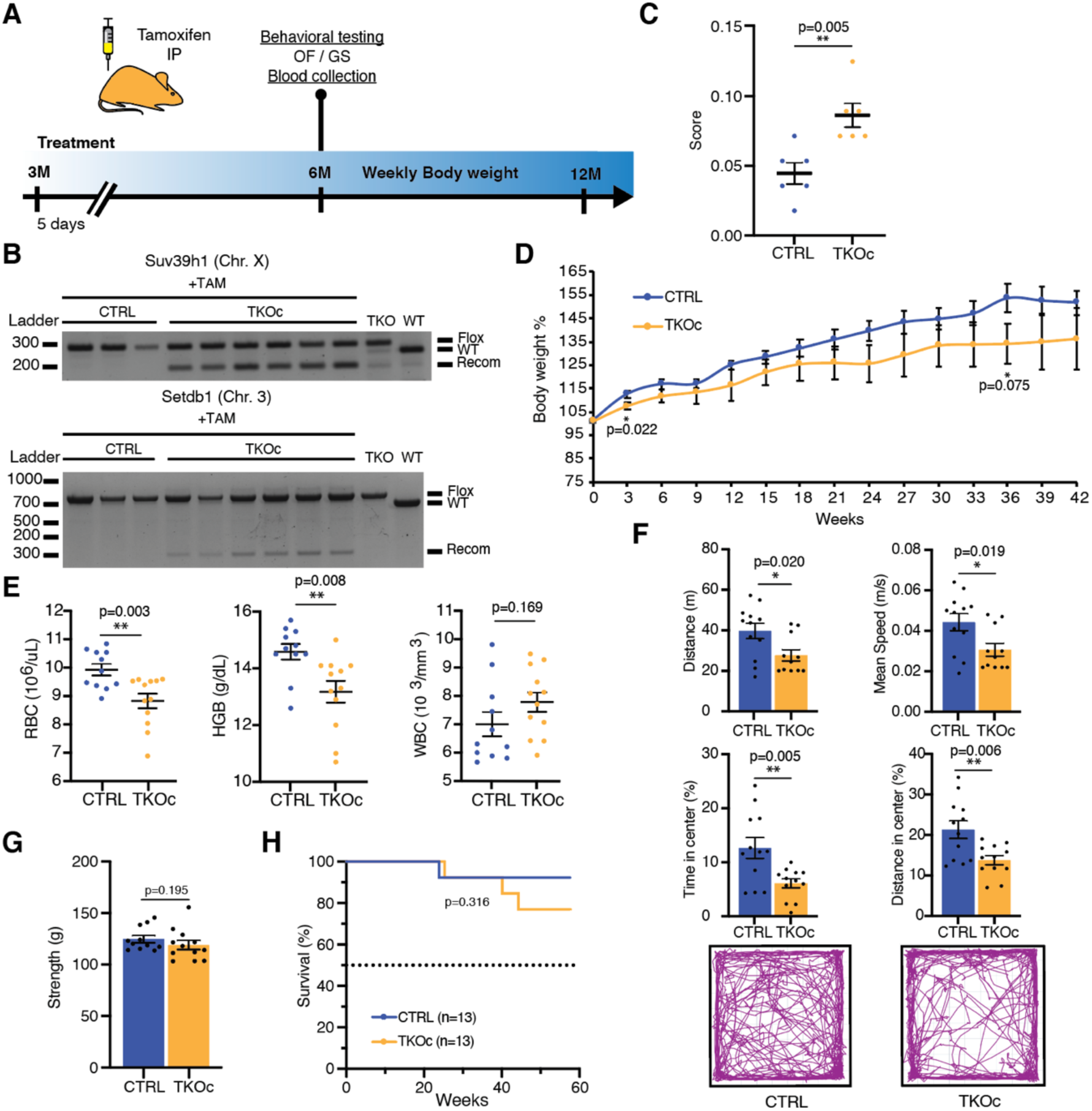
TKOc characterization upon single tamoxifen treatment. **(A)** Experimental design. **(B)** PCR analysis of Suv39h1 and Setdb1 recombination in blood from CTRL and TKOc mice upon Tamoxifen treatment. (n = 3 to 6). **(C)** Frailty index of CTRL and TKOc mice (n = 6 to 6, 6-month-old mice). **(D)** Changes in body weight of CTRL and TKOc mice, upon administration of tamoxifen. **(E)** Hematological parameters in CTRL and TKOc mice (n = 11 to 12, 6-month-old mice). **(F)** Behavioral characterization of CTRL and TKOc mice (6 months). Open field exploration (15 minutes), quantification of distance travelled, mean speed, time spent and distance traveled in the center with a representative tracking trace (n = 12 to 11-12, 6-month-old mice). **(G)** Grip strength measured as average grip strength of three different trials CTRL and TKOc mice. (n = 11 to 12, 6-month-old mice). **(H)** Kaplan-Meier survival curves for CTRL and TKOc mice (n =13 to 13 mice, including males and females). Dots in all panels represent individual sample data. Survival curve data were analyzed by log-rank (Mantel-Cox test). Data are mean ± SEM. Statistical significance was assessed by Two-tailed Student’s t test. *p < 0.05; **p < 0.01; ***p < 0.001.

**Figure S3:**
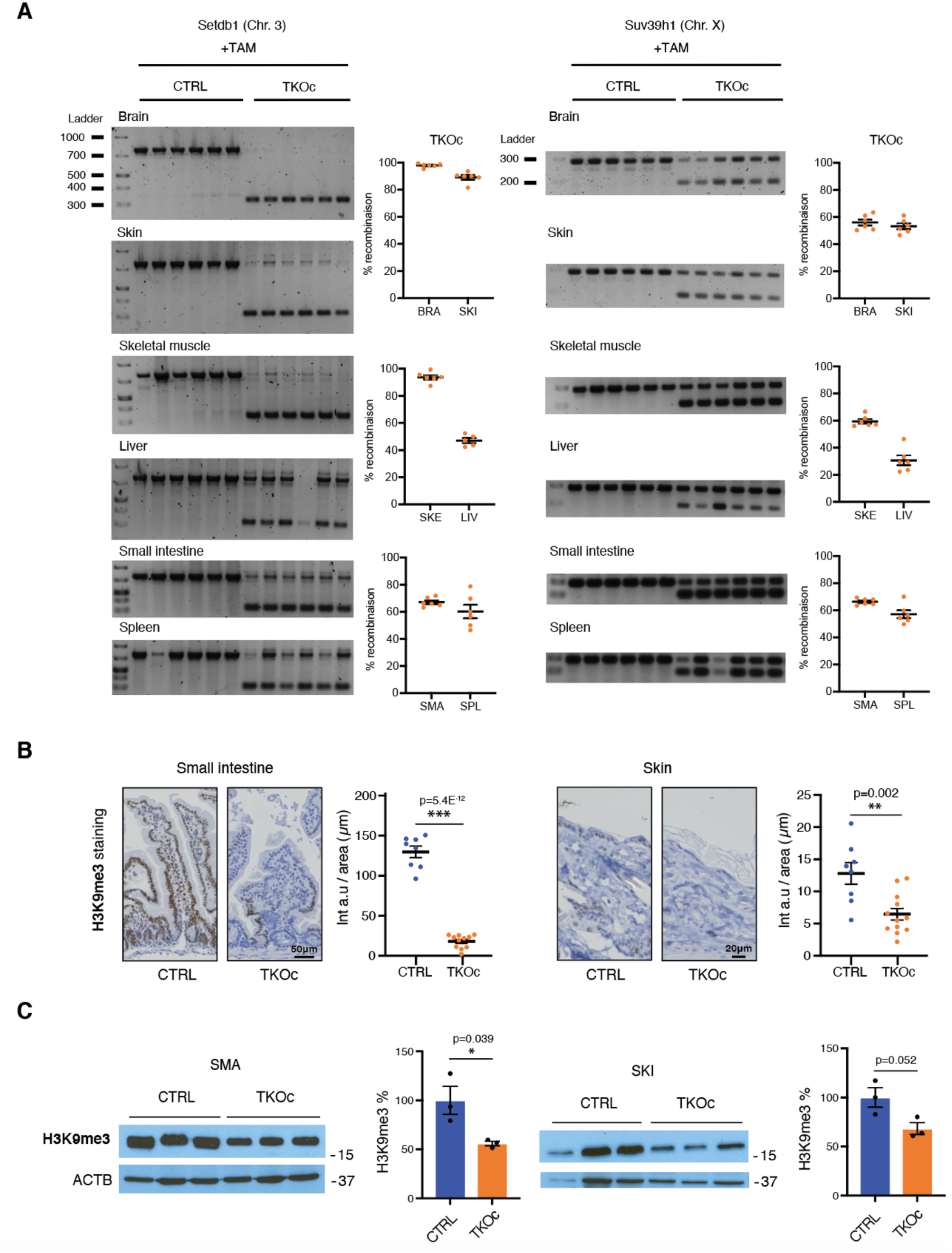
Analysis of recombination and H3K9me3 levels in TKOc and CTRL mice tissues. **(A)** PCR analysis of Suv39h1 and Setdb1 recombination in brain, skin, skeletal muscle, liver, small intestine and spleen from CTRL and TKOc mice upon Tamoxifen treatment. (n = 5 to 6, 7-8-month-old mice). **(B)** Immunostaining and quantification of H3K9me3 intensity in small intestine and skin of treated, CTRL and TKOc mice. (n = 2 to 3, 7-8-month-old mice). **(C)** Western blot of H3K9me3 in CTRL and TKOc small intestine and skin with TAM treatment. H3K9me3 quantification in CTRL and TKOc after treatment. (n = 3 to 3, 7-8-month-old mice). Data are mean ± SEM. Statistical significance was assessed by Two-tailed Student’s t test. *p < 0.05; **p < 0.01; ***p < 0.001.

**Figure S4:**
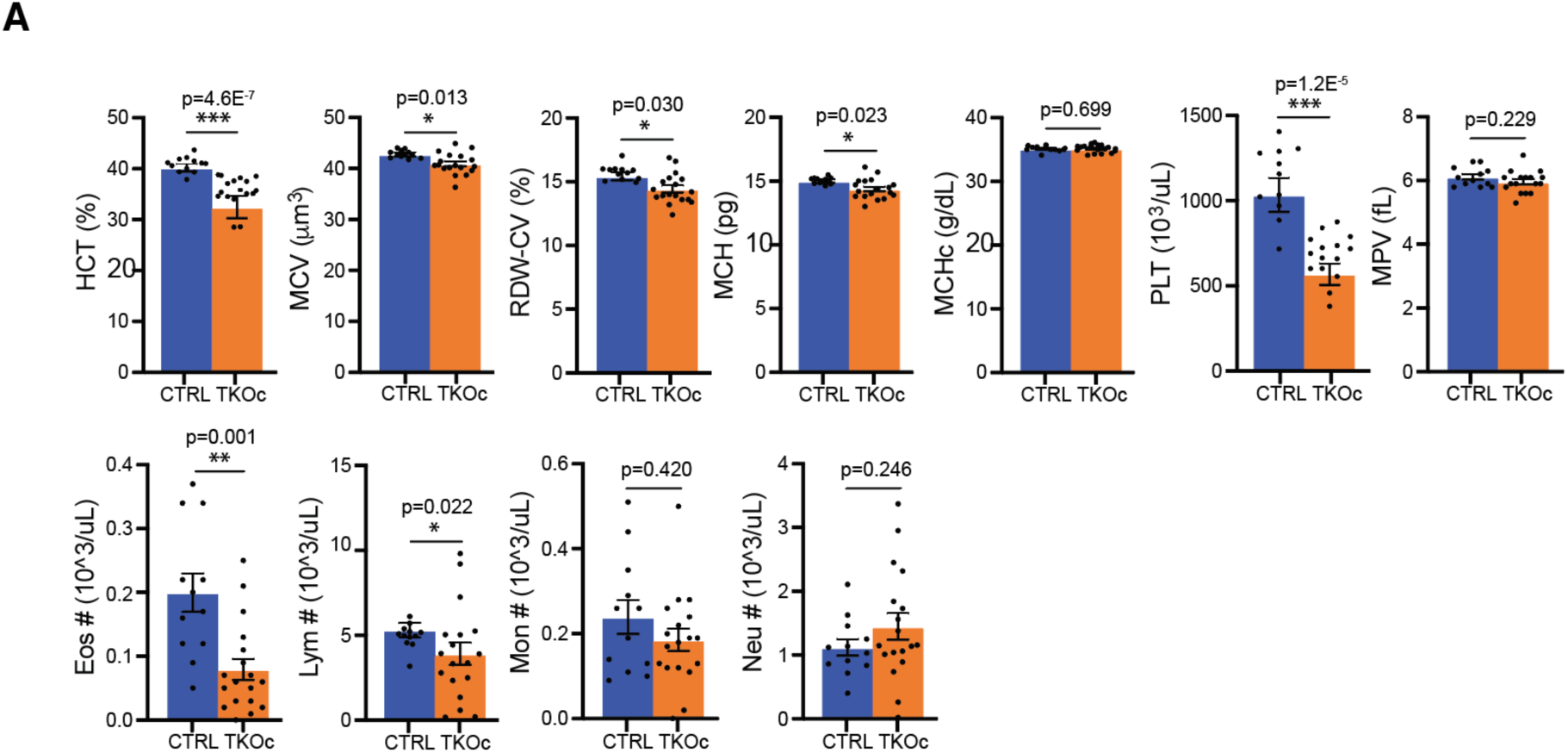
Analysis of blood from TKOc and CTRL mice. **(A)** Hematological parameters in CTRL and TKOc mice (n = 11 to 16, 6-month-old mice). Data are mean ± SEM. Statistical significance was assessed by Two-tailed Student’s t test *p < 0.05; **p < 0.01; ***p < 0.001.

**Figure S5:**
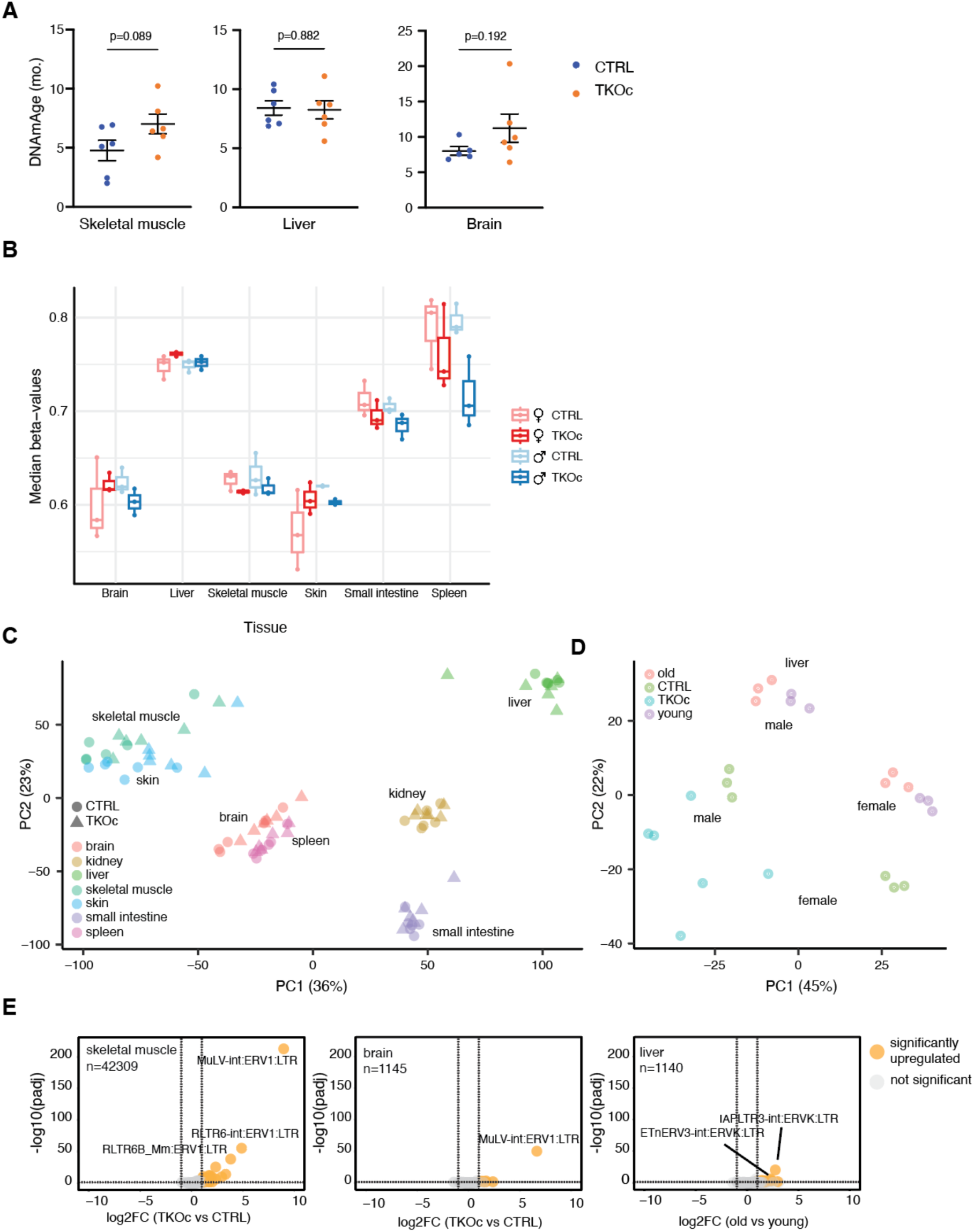
Comprehensive epigenetic and transcriptomic analysis from multiple tissues of TKOc and CTRL mice. **(A)** Epigenetic age of skeletal muscle, liver and brain of mice TKOc mice compare their CTRL (n = 6 to 6, 7-8-month-old mice). Data are mean ± SEM. Statistical significance was assessed by Two-tailed Student’s t test *p < 0.05; **p < 0.01; ***p < 0.001. **(B)** Overall DNA methylation levels estimated using the median beta-values. Each dot is a biological sample, and they are grouped by tissue, condition and sex. **(C)** PCA plot from total RNA-seq data derived from TKOc and CTRL mice tissues. **(D)** PCA plot from liver total RNA-seq data from TKOc, CTRL, young (3-months-old), and aged (18-months-old) C57BL/6JN mice. **e**, Volcano plots showing TE transcripts that are significantly upregulated (in orange), downregulated (in blue) or unchanged (in gray) in skeletal muscle, brain and liver from young and old WT mice. Transcripts were determined to be significant if the adjusted *p-*value was < 0.05 and log_2_fold change was > 1. A few top upregulated transcripts are labeled in each tissue.

**Table S1.**

Table showing the category of the top 50 significantly changed TE transcripts based on *p*-value in spleen, small intestine, skin, skeletal muscle, liver, kidney and liver from young and old WT mice

